# Exploring a mathematical framework for quantifying cell size-dependent glucose uptake in adipocytes

**DOI:** 10.64898/2026.02.26.707956

**Authors:** Christian Simonsson, Mathis Neuhaus, Jieyu Zhang, Karin G Stenkula

## Abstract

Insulin-stimulated glucose uptake (ISGU) in adipocytes is central to maintain systemic glucose homeostasis. Understanding how ISGU relates to adipocyte traits, such as cell-size, is critical for elucidating pressing questions related to metabolic dysfunction connected to adipose hypertrophy and hyperplasia. Cell size is considered a central trait reflecting multiple aspects of adipocyte metabolism. However, a robust quantitative approach to estimate ISGU for a specific cell size is currently missing. Here, in an attempt to move towards such a method, we have formulated an approach using a mathematical framework. The framework consists of a linear equation: the product of the known number of cells (calculated using coulter counter-based cell-size distributions) and the unknown ISGU/cell, compared to the absolute ISGU (measured using ^14^C-glucose tracer assays). To solve this equation, we formulate a minimization problem which is optimized to find the unknown ISGU/cell for the best solution. Using different formulations of the equation we can investigate the need for either cell size-dependent or independent ISGU/cell, to describe varying glucose uptake in a cell sample of various cell sizes. While the framework needs further refinement, we demonstrate that cell size-dependent uptake slightly improved the agreement between model and experimental data for some groups. Together, with further validation this could serve as a useful tool to resolve long-standing questions regarding size-dependent characteristics like adipocyte size and cellular function.

**Key findings:** Herein we explore a method to quantify cell size-dependent glucose uptake in adipocytes

## 1. Introduction

Sustained insulin-stimulated glucose uptake (ISGU) in adipocytes is essential to maintain systemic glucose homeostasis (1–3), and impairment of ISGU in in adipocytes is a driver of metabolic disease. Therefore, it is important to understand how different factors affect ISGU capacity in adipocytes. One of the determinants is the expression and translocation of GLUT4, the main insulin-responsive glucose transporter. Further, striking sex-specific differences related to ISGU capacity exists, where female mice display overall higher adipocyte ISGU than male mice (4). In a recent study, we monitored ISGU in adipocytes isolated from inguinal (ING) and perigonadal (PG) fat depots from female and male C57Bl/6J mice on regular chow diet (5), where we identified sex-specific and depot-specific differences. We also observed that glucose transport was related to mean cell-size for some groups (ING female, and PG male). A key question for understanding these differences is how ISGU capacity relates to adipocyte size.

Increased adipocyte size is amongst the most reliable predictors of metabolic dysfunction, supported by studies where subcutaneous adipocyte-size predicted insulin resistance independent of the degree of obesity (6–8). Since cell size varies greatly within the same fat depot, efforts have been made to characterize adipocyte function and ISGU in relation to size using various approaches. By single-cell imaging and cell filtration techniques, hypertrophic adipocytes were shown to display reduced cellular insulin sensitivity (9,10). However, manual cell sorting is cumbersome due to technical limitations resulting in inaccurate size separation, low yield and recovery of adipocytes and increased cell death. Therefore, ISGU is commonly assessed in relation to mean cell-size, comparing lean vs. obese subjects, or mice fed chow or HFD for various time (11–13). Together, while the presence of hypertrophic adipocytes correlates with impaired glucose homeostasis, their actual glucose transport capacity, and if there is a size-dependent cell-to-cell variation in the same fat depot, have scarcely been addressed. Thus, there is a need for alternative analysis methods.

Herein, we present a data-driven mathematical framework to estimate cell size-dependent glucose uptake for differently sized adipocytes isolated from C57BL/6J fed chow diet. The framework was used to analyse the presence of cell size-dependent glucose uptake in four different groups (male/female and ING/PG). We tentatively show that our approach could be used to quantify the glucose uptake for cells of different sizes, but more validation is needed.

## 2. Methods

In this study we used data from our previously published dataset of female and male C57BL/6J mice fed chow diet and terminated at different time points (5). The male group consisted of 30 mice, which were terminated at 12, 23, 35, 42, and 55 weeks of age. The female groups consisted of 24 mice that were terminated at either 12, 23, 35, and 42 weeks of age. All details regarding the glucose uptake measurement and coulter-counter based cell-size measurements can be found in the previous publication (5).

All mathematical and statistical analysis was done using MATLAB (MATLAB 2023b, MathWorks, Natick, Massachusetts). For all samples, we calculated the mean cell-size and the total number of cells based on the cell-size distribution data measured by coulter counter.

### 2.1 Mathematical framework for estimating glucose uptake

The following section will describe the model formulation and the parameter estimation process. Herein, we use two different model formulations to test two different hypotheses: either that the glucose uptake was size-independent, or size-dependent. The first model formulation was denoted M1 and was constructed for testing cell-size independent uptake, so that a single value of the ISGU/cell could be used for all adipocytes within a cohort (the same sex, and depot) to find agreement between the model and data. This can be written as a linear regression equation:

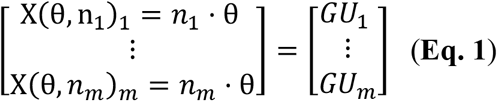

Each row represents an individual in the cohort, *n* is the number of cells in the 9µL solution used to measure the glucose uptake for an individual, *GU* denoted the total measured glucose uptake for the individual (mean value of technical triplicate in *fmol*), θ is the unknown model parameter representing the ISGU/cell, and *m* represents the last individual in the group (in male case 30, and in the female case 24). The number of cells was calculated using coulter counter-based measurement of cell-size distribution. Importantly, note that the θ is the same for all rows. The second formulation M2 was constructed to test cell size-dependent uptake and dictates that the ISGU/cell should change depending on adipocyte-size. Here, different cut-off intervals based on the cell diameter were used to subdivide the total number of cells. For the results presented herein we choose to divide the distribution into three adipocyte subgroups based on the cell diameter (Ø): Ø < 70 µm, 70µm ≤Ø≤ 100 µm, 100µm < Ø. Based on the subgroups we can subdivide the regression equation of glucose uptake by modifying **Eq. 1**:

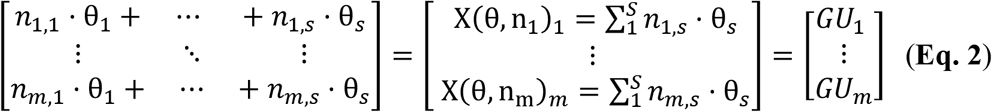

where θ_1_ to θ_*s*_ is are the parameters dictating the ISGU/cell for each cell size group and where each cell size group has an index 1 to *s*. To also account for group (age) difference we added group specific parameters that allowed changed ISGU/cell, this formulation was denoted M3 and M4. Where M3 doesn’t have any cell size-dependent uptake. The equation for M3 can be written as:

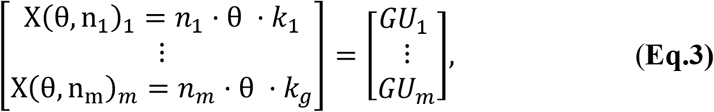

and for the M4 formulation it can be written as:

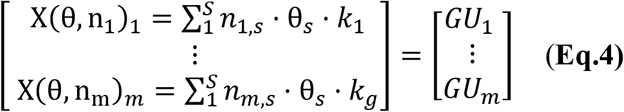

where *k* are the free parameters representing the change in glucose uptake related the group (*g*). Moreover, in all model formulations to estimate the unknown parameters (θ, or the set θ_1_ to θ_*s*_) we minimize the difference between the model estimated glucose uptake and the measured data. We also formulated models where the ISGU/cell instead was dependent on size using a continuous function. This was formulated without and group-specific parameters. By reformulating **Eq. 2** we get the following equation:

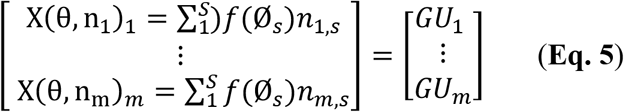

where *f* is a continuous function describing the ISGU/cell relationship to the cell size diameter Ø, where *S* is the last bin, starting from bin size 20um to bin size 240um with 0.5 um increments. Using this formulation, we could systematically test different functions *f* to compare different behaviors i.e. only allow for increasing or decreasing ISGU/cell with increasing cell diameter. We tested four different formulations: M5 a linear relationship between cell size and ISGU/cell (one free parameter); M6 a gaussian curve (three free parameters); M7 is a negative linear relationship (one free parameter); and M8 is a log-linear relationship (one free parameter).

For all model formulation we performed parameter estimation. The parameter estimation process can be described using the following minimization problem. In practice, this was done by reformulating the equations (either **Eq. 1** for the M1 case, **Eq. 2** for the M2, **Eq. 3** for the M3 and **Eq. 4** for the M4 case, **Eq.5** for the rest) into a minimization problem which can be written as:

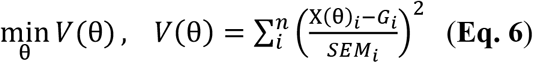

where *V*(θ) is the objective function, which was minimized with respect to the unknown parameters (different ISGU/cell), X(θ)_*i*_ was the model estimated glucose uptake for mouse *i*, and the *SEM*_*i*_.was the standard error of the mean (SEM). The SEM value was set to be 20% of the mean value for each data-point. The minimization was performed using the MEIGO toolbox (14) in MATLAB. The agreement between model and measured glucose uptake was evaluated using a χ^2^-test on the residuals, where we used a 5% confidence level and the number of data points as the degree of freedom.

## 3. Results

To resolve how glucose uptake varies depending on cell size, we investigated the relationship between the estimated mean cell-size (in samples with a fixed sample volume, see methods) and the corresponding ISGU/cell. We hypothesized that two scenarios exist: cell-size independent uptake or cell-sized dependent uptake (see methods section **Eq. 1** and **Eq. 2**). To explore this further we investigated the relationship between glucose uptake and cell size and used a mathematical framework to see if size-dependent uptake can be quantified.

### 3.1 Insulin-stimulated glucose uptake compared to mean cell-size

To investigate these two scenarios, based on our previous findings (5), we performed linear regression analysis between the mean cell-size with ISGU/cell for each depot in respective sex. For the ING depot in males, no correlation was found between mean cell-size and ISGU/cell (R^2^=0.01) (**Fig. 1A**). In contrast, for the ING female depot, we found a positive correlation between ISGU/cell and mean cell-size (R^2^=0.63) (**Fig. 1B**). This indicates that different sized cells have different ISGU in the ING female depot. For the male PG depot, a positive trend (R^2^=0.45) between mean cell-size and ISGU/cell (**Fig. 1C)** was observed, and in the PG female depot we observed no correlation between mean cell-size and the ISGU/cell (R^2^=0) (**Fig. 1D**). The relationships observed in the ING female and PG male support the notion that adipocytes of varying sizes have different glucose uptake capacity in response to insulin.

**Figure 1:**
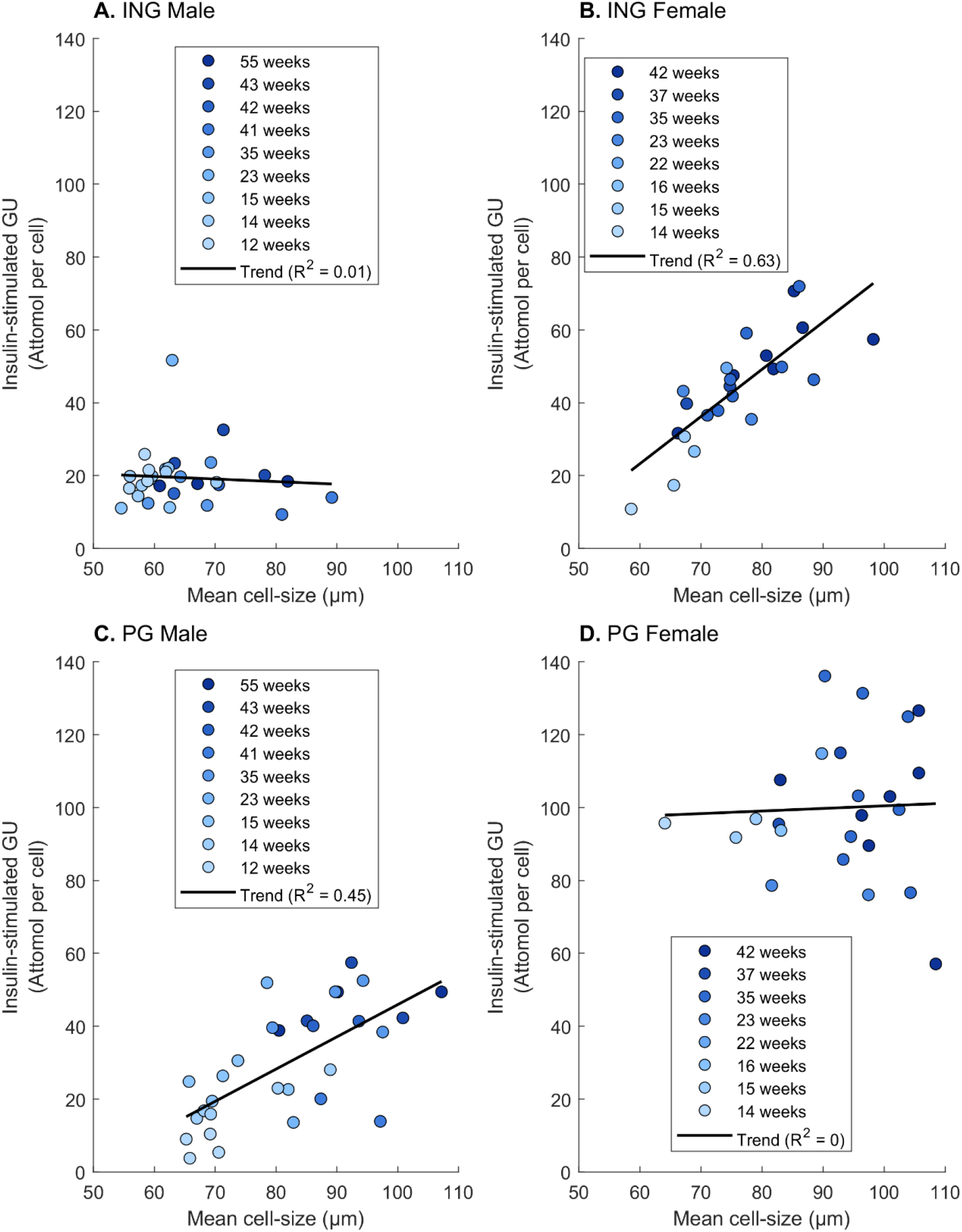
Regression analysis between insulin stimulated glucose uptake per cell (ISGU/cell) and the mean cell-size. **(A).** For the ING depot male, **(B)** ING depot female, **(C)** PG depot male, and **(D)** PG depot female.

### 3.2 Introducing an analysis framework for estimating size-dependent ISGU

Next to further elucidate this relationship, we created a mathematical framework, depicted in **Fig. 2**. The framework can be described by several steps. Briefly, in the first step (**Fig.2A**), data is collected on cell-size distribution (**Fig. S1**), total cell number and absolute ISGU (in a fixed sample volume) for a population of mice. In the second step (**Fig.2B**) the data is integrated into a (regression) equation representing either of the two scenarios M1 or M2 (see methods section **Eq. 1** and **Eq. 2**). In the third step (**Fig.2C**), the model parameters representing the unknown ISGU/cell (θ, or the set θ_1_ to θ_*s*_) were estimated by using an optimization algorithm to find the parameter values resulting in the smallest difference between model and measured data. In the last step (**Fig.2D**), we evaluated the agreement between the approximated solution and measured glucose uptake using a χ^2^ test of the residuals.

**Figure 2:**
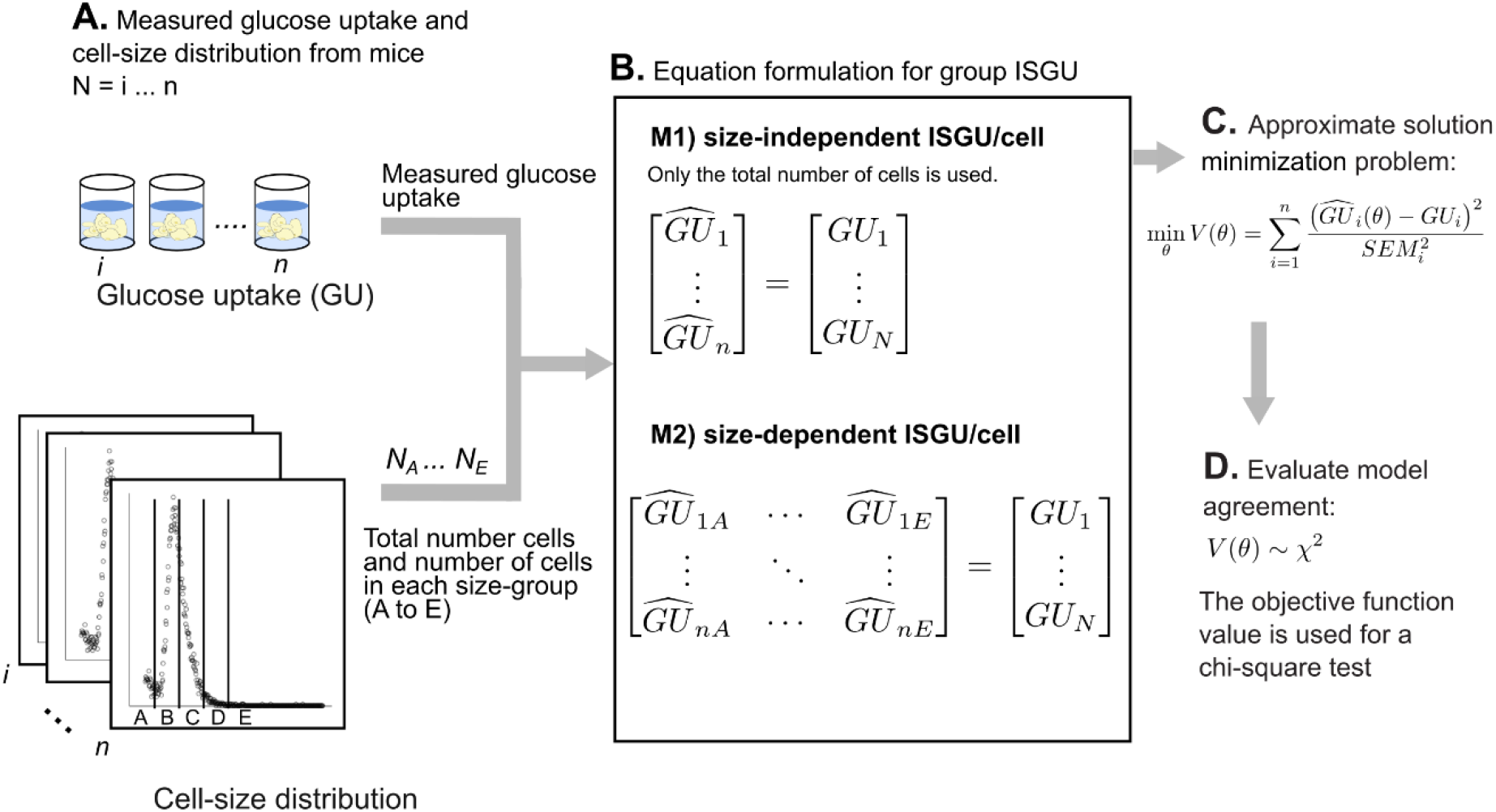
Schematic of the mathematical framework. The framework consists of several steps. In the first step **(A)**, measurements of the glucose uptake and the cell size distribution were collected from each mice in the cohort N, consisting of mice with indexes i ton. In step **(B)** the cell size distribution measurements (and the number of cells) were integrated into a mathematical expression describing either M1) cell-size independent ISGU/cell or M2) cell size-dependent ISGU/cell. This expression was used as the left-hand side as part of an equation equaling the measured glucose uptake. **(C)**. In the third step we tried to solve this equation by estimating value of the unknown parameters θ, or the set θ_1_ to θ_s_. **(D)**. In the last step, the agreement between the approximated solution and measured total glucose uptake was compared using a χ^2^-test on the residuals.

As an initial step we used the framework to estimate the ISGU/cell within each group (based on age, sex, depot) separately (**Supplementary Table S1**). For this analysis we observed that the M1 formulation was sufficient to pass the χ^2^-test in most cases (not PG male 12 weeks and 23 weeks). Running the analysis with the M2 formulation produces a slight improvement in the cost-value (Supplementary **Table 1**). Performing the analysis for each group separately is limited by the fact that same aged individuals have similar number of cells (similar cell size distributions) and is thus not ideal for our purpose to explore quantification of cell size dependent uptake.

Thus, to test if the framework could be used to quantify the cell size-dependent uptake we pooled data from the same sex and depot into groups. As observed in **Fig. 1**, we know that for the PG female and ING male the ISGU/cell could be more size-independent, as no correlation was observed between ISGU/cell and size. As expected, the M1 formulation could produce a qualitatively good agreement to the pooled data for these cases (**Fig. S2A-B**). For the ING male case the model fails to describe the data for several mice most notably for 2, 14 and 16 (**Fig. S2A, blue curve**). Moreover, for the other two cases, the ING female and PG male, where we observed a size-dependent ISGU/cell in Figure. 1, we observed moderate improvement using the M2 formulation (**Fig. 3A-B**). For the ING female case (**Fig. 3A**) the M2 model reduce the cost-value from 158.6 to 50.3, but the agreement does not pass a χ^2^-test, as well as fail to describe qualitative difference in mice 1 to 4. For the PG male case the M2 improvement is not noticeable qualitatively, with a reduction from 365.1 to 241.1 in cost value. To better describe these data, we next investigated how to account for group effects.

**Figure 3:**
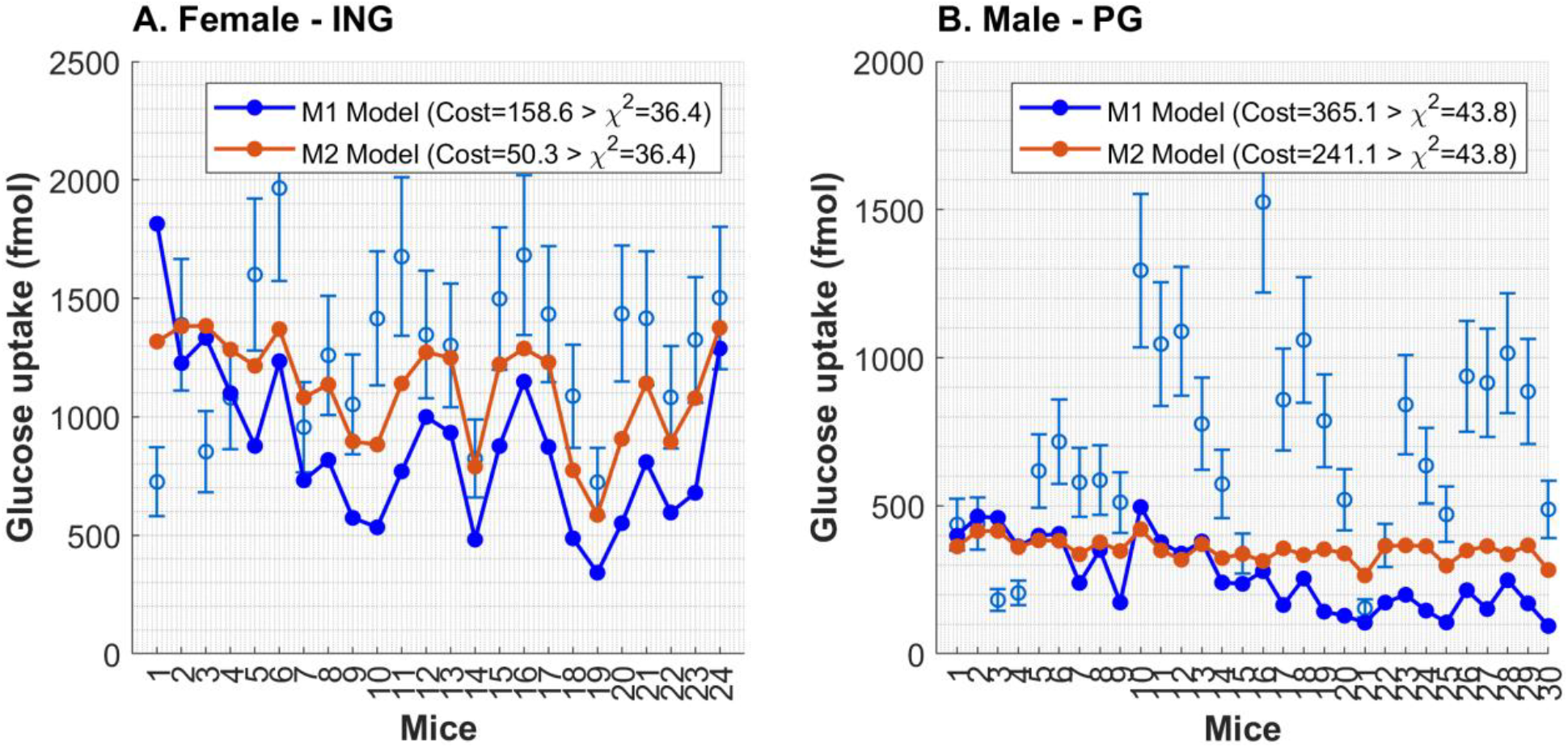
Model agreement with pooled data. Two model formulations were tested: M1; same ISGU/cell, M2; size-dependent ISGU/cell **(A)**. Agreement for the female ING depot. **(B)**. Agreement for the male PG depot.

### 3.3 Evaluating quantification of cell size-dependent glucose uptake

To further improve the agreement, we tested the M3 and M4 formulation (**Eq. 3** and **Eq. 4**) which allows for group (age)-specific differences in the ISGU/cell (see Methods section). For the ING female case both the M3 and M4 model passes the χ^2^-test, with a cost value of 22.9 and 11.4 respectively (Tχ^2^ = 36.4) (**Fig. 4A**). Still, some mice could not be explained, for example mouse number 2. For the PG male a large improvement can be seen and both the M3 and M4 model passes the x2-test, with a cost value of 42.1 and 37.7 respectively (Tχ^2^ = 36.4) (**Fig. 4B**). Also, here the ISGU for some mice 1, 2 and 16 cannot be described fully. Because the M3 and M4 pass the test in both cases, it is not obvious if the cell size-dependent uptake is true. For the ING female the M4 model has a larger reduction in cost, and cell size-dependent uptake could maybe be argued for in this case. Nevertheless, the solution for the M4 formulation for both cases is presented in Figure 4C-D. Here, we can see that the model fitting in both cases choses the smaller cell (Ø < 70 um) to take up the least per cell. However, due to the fact the M3 formulation produces a similar acceptable solution, one cannot disregard that the difference in ISGU/cell can be due to age, or group related effects such as insulin resistance, or other natural variations, rather than explicit difference in glucose uptake related to cell-size.

**Figure 4:**
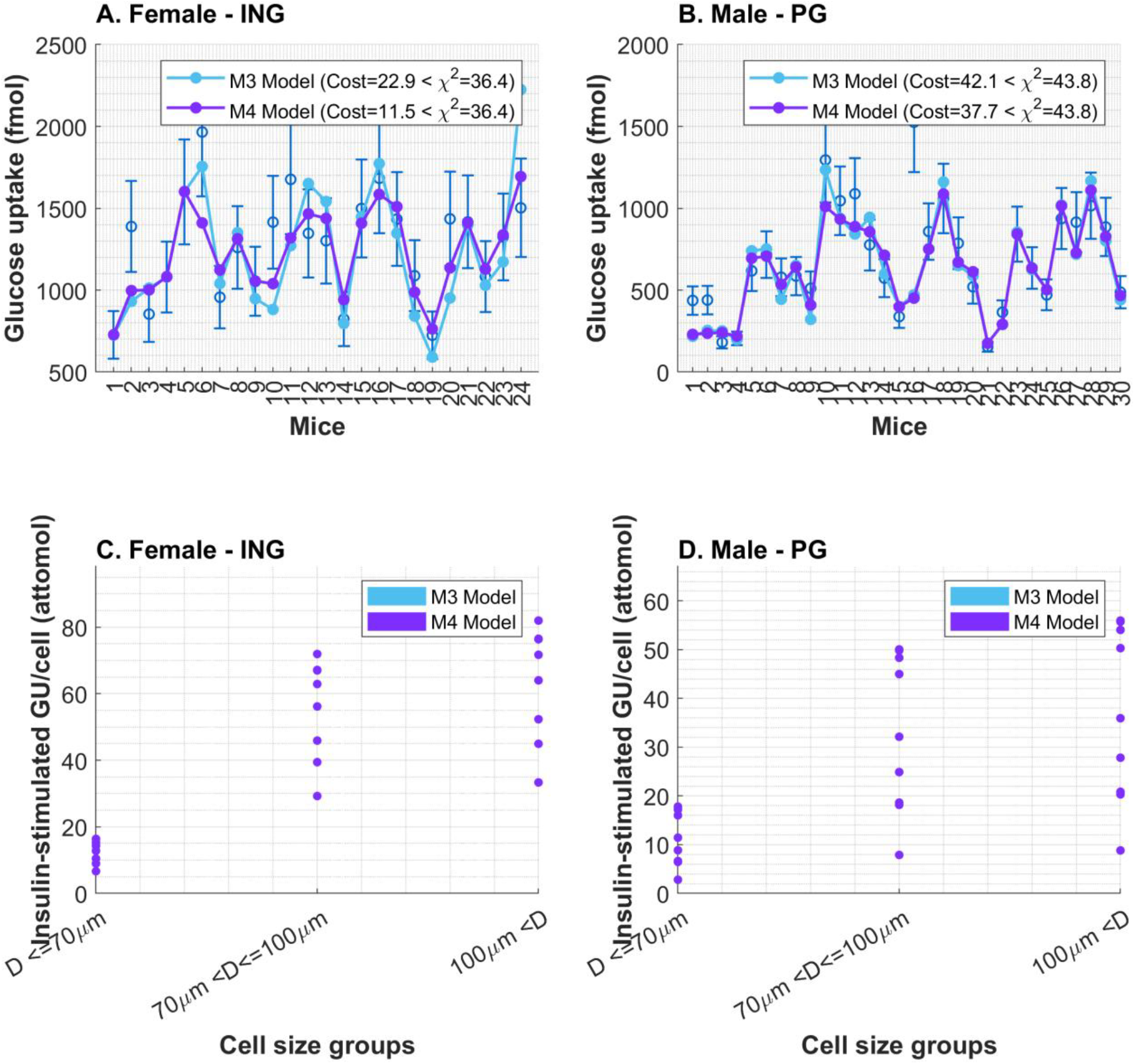
Model agreement with pooled data when for group-group differences. Two model formulations were tested: M3; same ISGU/cell, M4; size-dependent ISGU/cell, both with group-group difference allowed. **(A)**. Agreement for the female ING depot. **(B)**. Agreement for the male PG depot. **(C)**. Model estimated ISGU/cell for the M4 model formulation for the female ING depot. **(D)**. Model estimated ISGU/cell for the M4 model formulation for the male PG depot.

### 3.4 The framework can be used for hypothesis testing

We also tested an approach where the ISGU is expressed as a continuous function dependent on cell-size (**Eq. 5**). Using this approach, we could investigate different relationships between cell-size and glucose uptake. Here we show this approach as an illustrative example using ISGU from the ING female group (**Fig. 5**). We tested four different formulations: M5 describes a linear relationship between cell size and ISGU per cell; M6 represents a Gaussian curve; M7 describes a negative linear relationship; and M8 represents a log-linear relationship. As can been seen there is no improvement in cost-value for the M7 formulation, but a slight improvement for the others, with M6 having the largest decrease (**Fig. 5A**). For each formulation, the corresponding continuous ISGU/cell compared to cell size can be observed (**Fig. 5B**).

**Figure 5:**
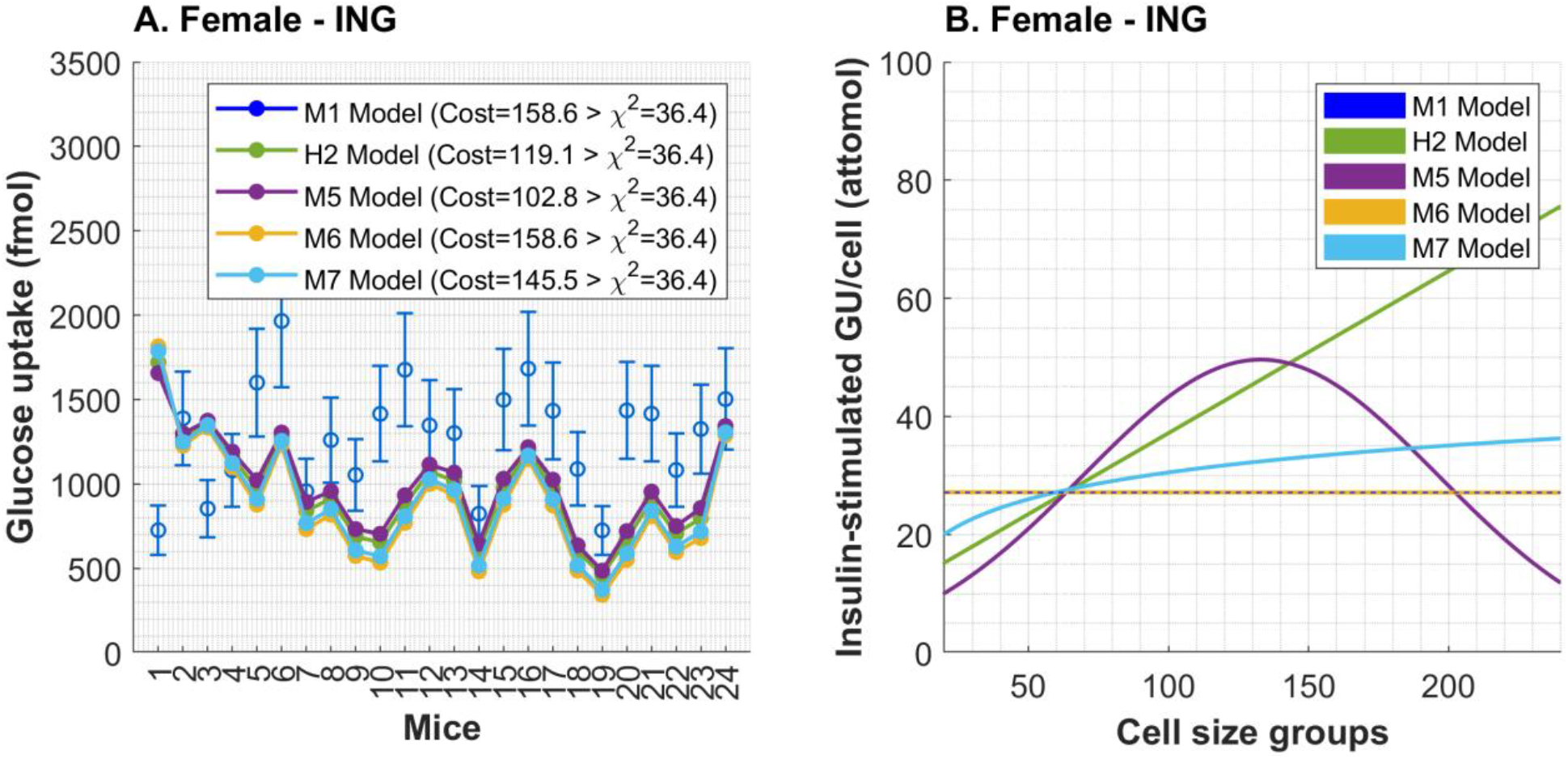
Model agreement using continuous function for cell size-dependent glucose uptake. Here, we tested four model formulations with different functions describing the relationship between ISGU/cell and cell diameter. We tested four different formulations: M5 describes a linear relationship between cell size and ISGU per cell; M6 represents a Gaussian curve; M7 describes a negative linear relationship; and M8 represents a log-linear relationship. **(A)**. Model agreement between all formulations, including M1 (for comparison) for the ING female depot **(B)**. The found solution after parameter estimation, showing the shape of the ISGU/cell relationship to cell diameter for the different formulation. No solution passes a χ2-test.

## 4. Discussion

Herein we present a mathematical framework for investigating ISGU in adipocytes from male and female C57BL/6J mice fed chow diet. We have previously (5) discovered that adipocytes isolated from female PG and male ING fat depots had comparable ISGU/cell independent of size, while male PG and female ING displayed varying ISGU/cell in a cell size-dependent fashion (**Fig. 1**). This was further investigated herein, using a mathematical framework (**Fig. 2**). We tested two main scenarios using this approach, M1 and M2. In scenario M2, which allowed the ISGU/cell to depend on cell size, it was possible to improve the fit for ING females, but this was not sufficient to fully describe the data and resulted in only a slight improvement for PG males (**Fig. 3**).To test if a group-specific effect is needed to describe data in these two cases – we introduce model formulations M3 and M4. We showed that both the M3 and M4 formulations could describe data for both ING female and PG male, meaning that the difference in glucose uptake could be due to natural or technical group-group difference (**Fig. 4**). Lastly, we also show how the framework could be used to evaluate different relationships between ISGU/cell and cell diameter (**Fig. 5**). Nevertheless, for the ING female the M4 formulation produced a substantial improvement in agreement. However, further validation and model development to understand sources of variance is still required.

Reduced ISGU in hypertrophic adipocytes under obese condition is well known (15). In contrast, not many studies have investigated difference in cell-size related difference in ISGU during lean conditions. There are several studies making use of buoyancy separation for studying cell-size related differences in adipocytes. In a study by Skruk et al (16), adipokine expression is shown to differ in four different cell-size groups. Similar methods have been used to measure size-dependent lipolysis (17), and gene expression (18) in adipocyte. However, studies measuring glucose uptake in size-separated adipocytes are sparse (especially in lean condition). In future work, it would be valuable to investigate whether combining our mathematical framework with data from flotation-sorted cell fractions improves performance in quantifying cell-size-dependent uptake.

We investigated cell size-dependent ISGU using our presented framework. Consistent with our previous study (5), and based on the same data, the PG male depot behaves similarly to the ING female depot, exhibiting a correlation between ISGU per cell and mean cell-size (R^2^ = 0.45; Fig. 1C). For this group, the M1 formulation did not find a solution (**Fig. 3**), neither could the M2 formulation. Nevertheless, introducing the M2 formulation improved the agreement in the ING female case, which could be indicative that cell size-dependent glucose transport capacity could explain some of the variability in data. However, none of the model formulation passes the χ^2^ test for any combination of data, and the PG male case does neither fit key qualitative behaviors when using M1 or M2. The M3 and M4 produce a sufficient solution, which is indicative of that the group-group or age effect is larger contributor for explaining the variance in data. Also, no increasing or decreasing trend is observed for ISGU in these groups i.e., that ISGU would increase with age. This might be indicative of the fact that the variance between groups is either due to natural (insulin sensitivity) or technical variations (cage-cage effect, stress) between the groups. The current model does not account for any subject-subject specific variability beyond the explainability of e.g. mean cell-size and depot weight. Our analysis indicates that the cell size-dependent ISGU/cell displays distinct depot-specific behaviors in females; either constant ISGU/cell (PG) or increased ISGU/cell (ING) in larger cells. In male mice, this depot-specific distinction is not as clear.

There are several aspects of the framework and the analysis that should be discussed. The groups we analyzed were divided based on sex and fat depot. Consequently, we ran the analysis separately for each group and therefore did not have to account for sex- or depot-specific differences in the model formulation. Still, we also analyzed the predictive effect of multiple different variables (**Fig. S3**). Interestingly, as also observed in Figure. 1, and previously (5), we saw opposite behaviors between depots in the female groups. This might indicate that in females, adipocyte glucose uptake is not sensitive to changes in e.g. body or depot weight. Moreover, the male depot had more similarities in their relationship with these parameters. Nevertheless, we tried to add these variables (individually) to the model equations, but they did, as we expected, not add any explainability. One could speculate that this is due to the fact that body- and depot-weight already are adjusted for in the number of cells for ING male and PG female, and thus no additional explainability is added to the regression model. For the model to more accurately describe the variance in data, and to increase the predictive capability, more work needs to be done to incorporate factors beyond cell-size that could impact inter-subject variance.

Another aspect of the model formulation is the number of cell-size groups, which was limited by the number of size bins used in the coulter counter measurements (in our case 400 bins ranging from 20µm to 250µm, with 0.5 µm increments). The model can be formulated to include all cell-size (by expanding number of columns in **Eq. 2**). Such implementation will increase the model complexity, as a single parameter is used to depict the ISGU/cell for each cell-size. This implementation would require more data to parameterize; and thus, we currently choose a minimal approach of only using three cell-size groups. Alternatively, we herein also demonstrate that one can use a continuous function dependent on the cell size (diameter) for describing ISGU/cell for each bin size (**Fig. 5**). This approach could also be a tool to evaluate hypothesis regarding certain behaviors *i*.*e*. that the ISGU/cell should increase or decrease with size. This approach is limited by the choice of functions, compared to using free parameters for different size-groups. Importantly, the framework presented herein is not limited to glucose uptake, but could be used to estimate any cellular function, e.g. lipolysis which is linked to cell-size in healthy subjects (17). Additionally, our method could also make use of microscopy data on cell-size distribution (with corresponding glucose uptake data).

There are not many studies that look into quantification of uptake related to cell-size, especially in primary cells, as we do herein. Nevertheless, a study by Khetan et al. (19) highlights that under both reaction- and diffusion-controlled regimes, cellular uptake follows a linear relationship with the cell radius. Interestingly our observation in adipocytes indicates that this is unlikely in the case of ING male and PG female (Fig. 1B and Fig. 1D) but would be a more likely case for ING female. However, our findings need further validation.

There are some inherent limitations with our framework, one being that the analysis cannot be done on a single sample. Rather it is a population-based regression analysis and alas to estimate the cell size-dependent ISGU/cell there needs to exist a difference in the cell-size distribution (within the population). Another limitation is the used noise-level, as the data is triplicate ISGU measurements from individuals, we have to assume a standard deviation of the mean as we applied a χ^2^ test. Here we assumed 20 %, which might be too low in some cases, if compared to the actual SEM calculated for a group. However, as we wanted to describe the difference between individuals and having a cut-off for the goodness of fit, we used the current approach. Alternatively, one could skip the noise-level assumption and evaluate the qualitative agreement to the data or regression measures such the coefficient of determination (R^2^).

There also are some limitations for the data used. Adipocyte size was measured in isolated adipocytes, and very small adipocytes are likely removed during the washing process due to their low buoyance. Also, the normalization of our readouts to the cell number could be influenced by differences in the packing of cells after floatation and the abundance of small cells. We assume that the effects of this are minor since we ensured constant floatation times and precise handling.

Together, we have investigated the relationship between insulin-stimulated glucose uptake capacity (ISGU/cell) and adipocyte size. We found that cell size-dependent ISGU/cell could improve the agreement between model and experimental data although only slightly. Nevertheless, this serves as a first attempt towards an analysis framework for quantifying adipocyte function in relation to size. With further validation and development, our approach could serve as a tool to generate novel insights, elucidating the relationship between adipocyte size and cellular function.

## Acknowledgements

Not applicable.

## Author Contributions

CS, MN, JZ and KS confirm contribution to the paper as follows: study conception and design: CS and KS. Analysis and interpretation of results, draft and critical review of manuscript: CS, MN, JZ and KS. All authors reviewed the results and approved the final version of the manuscript.

## Ethics approval and consent to participate

Not applicable.

## Availability of data and materials

Not applicable.

## Supplementary

**Figure S1:**
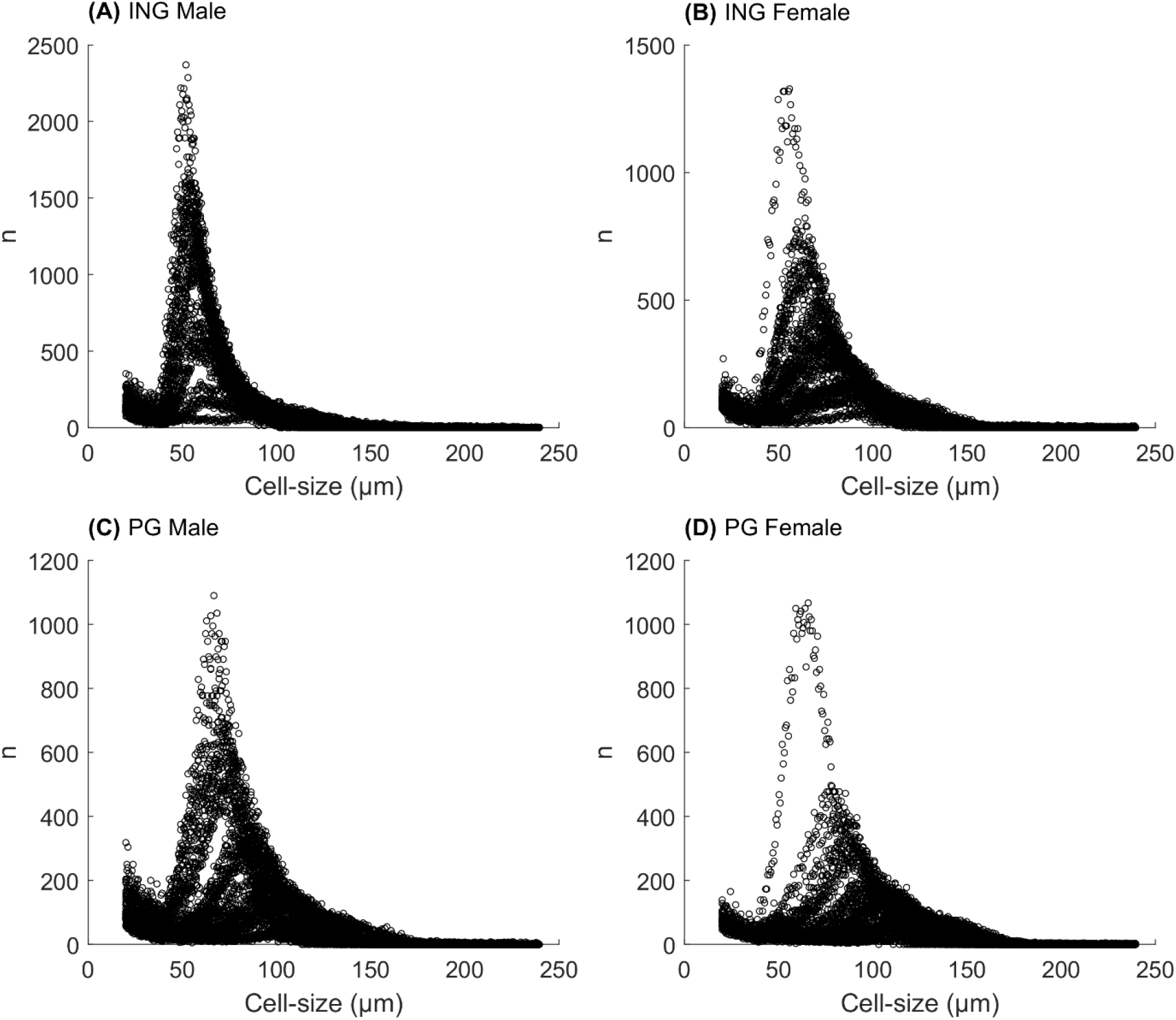
Cell-size distribution measured by coulter counter. **(A)**. Distribution for the male Inguinal (ING) depot. **(B)**. Distribution for the female ING depot. **(C)**. Distribution for the male Perigonadal (PG) depot. **(D)**. Distribution for the female PG depot.

**Table S1:** The cost-value (χ^2^ test statistic) for the agreement between model formulation (M1 or M2) when compared to training data for each group based on depot, sex and age. The χ^2^ test limit (Tχ^2^) using confidence level, α=0.05, and degree of freedom equal to number of data points for each group is presented in the last column.

| Depot | Sex | Group (Age in weeks) | Cost (M1) | Cost(M2) | $T\chi^2$ (limit) |
| --- | --- | --- | --- | --- | --- |
| PG | F | 14 | 0.0* | 0.0 | 3.8 |
| PG | F | 15 | 0.0 | 0.0 | 6.0 |
| PG | F | 16 | 0.0 | 0.0 | 3.8 |
| PG | F | 22 | 0.0 | 0.0 | 3.8 |
| PG | F | 23 | 0.0 | 0.0 | 6.0 |
| PG | F | 35 | 7.4 | 6.7 | 15.5 |
| PG | F | 37 | 0.4 | 0.0 | 6.0 |
| PG | F | 42 | 12.2 | 7.6 | 14.1 |
| ING | F | 14 | 0.0 | 0.0 | 3.8 |
| ING | F | 15 | 3.6 | 2.7 | 6.0 |
| ING | F | 16 | 0.0 | 0.0 | 3.8 |
| ING | F | 22 | 0.0 | 0.0 | 3.8 |
| ING | F | 23 | 0.5 | 0.0 | 6.0 |
| ING | F | 35 | 7.6 | 4.0 | 15.5 |
| ING | F | 37 | 0.2 | 0.0 | 6.0 |
| ING | F | 42 | 11.1 | 1.8 | 14.1 |
| PG | M | 12 | 14.4 | 13.3 | 9.5 |
| PG | M | 14 | 6.2 | 0.4 | 11.1 |
| PG | M | 15 | 2.8 | 1.9 | 11.1 |
| PG | M | 23 | 12.8 | 11.8 | 6.0 |
| PG | M | 35 | 1.8 | 1.6 | 9.5 |
| PG | M | 41 | 1.6 | 0.0 | 6.0 |
| PG | M | 42 | 0.0 | 0.0 | 6.0 |
| PG | M | 43 | 1.5 | 1.3 | 7.8 |
| PG | M | 55 | 1.0 | 0.1 | 7.8 |
| ING | M | 12 | 2.6 | 1.7 | 9.5 |
| ING | M | 14 | 2.2 | 2.0 | 11.1 |
| ING | M | 15 | 10.5 | 8.6 | 11.1 |
| ING | M | 23 | 7.1 | 6.7 | 6.0 |
| ING | M | 35 | 7.4 | 6.8 | 9.5 |
| ING | M | 41 | 1.9 | 0.5 | 6.0 |
| ING | M | 42 | 0.3 | 0.0 | 6.0 |
| ING | M | 43 | 2.6 | 2.0 | 7.8 |
| ING | M | 55 | 0.1 | 0.0 | 7.8 |

\*Smaller than 0.1

**Figure S2:**
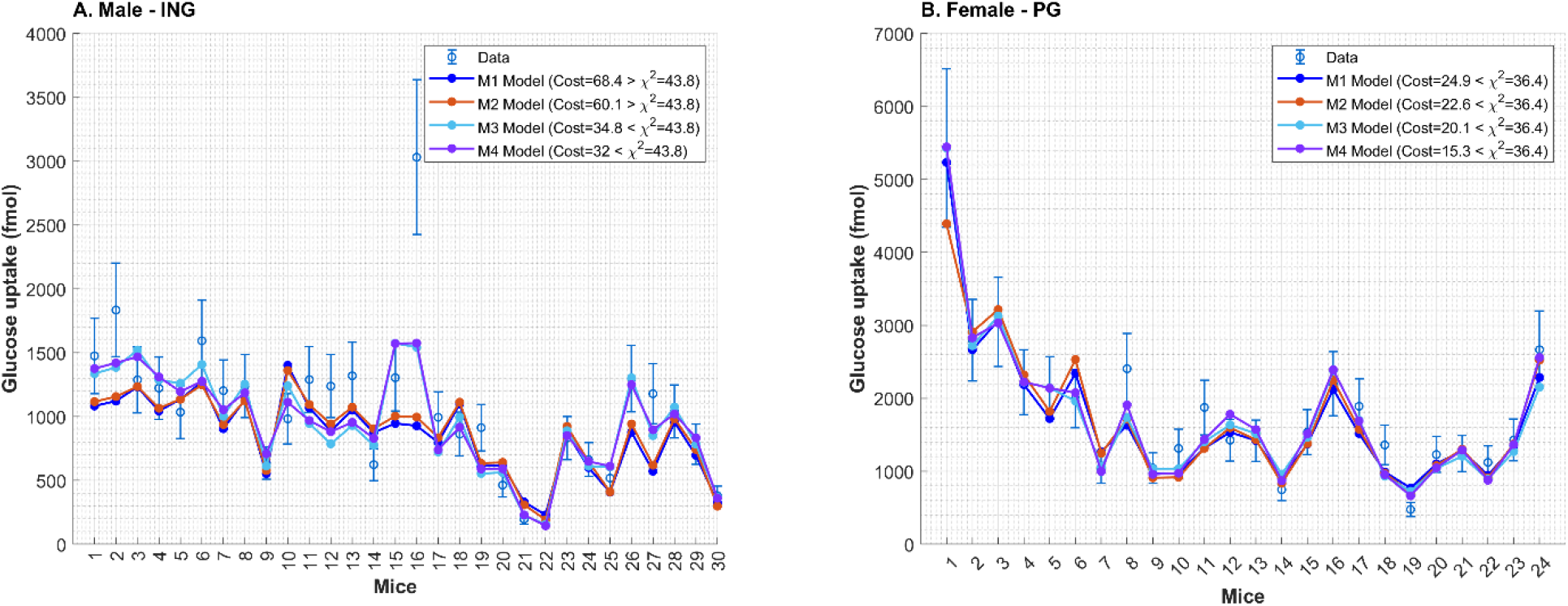
Three model formulations were tested: M1; same ISGU/cell. M2; size-dependent ISGU/cell and M3 size-dependent ISGU/cell with age-age variance. **(A).** Agreement for the male ING depot. **(B)**. Agreement for the female PG depot. **(C)**. Model estimated ISGU/cell for the three different model formulations for the male ING depot **(D)**. Model estimated ISGU/cell for the three different model formulations for the female PG depot

**Figure S3:**
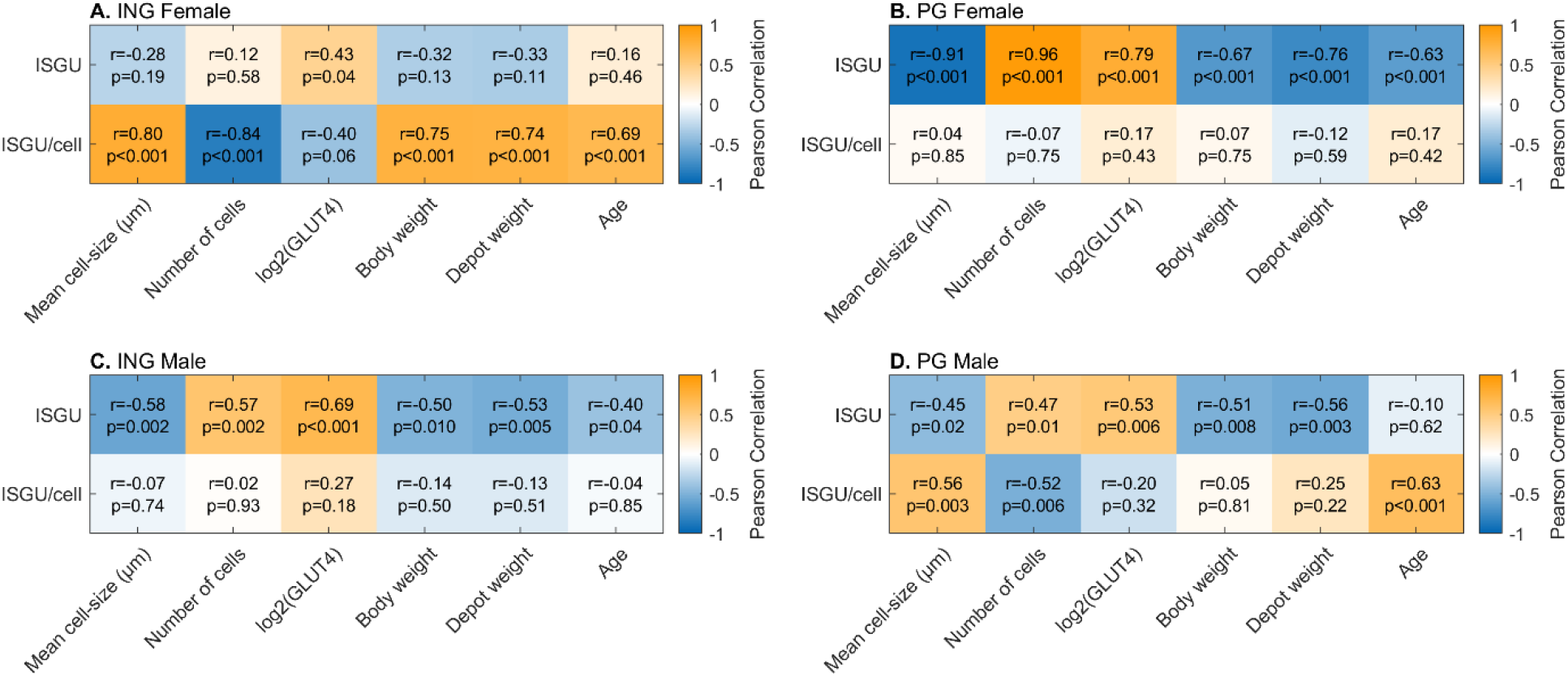
Pearson correlation matrix showing possible covariate relationships for glucose uptake. **(A).** Correlation matrix for female ING glucose uptake. **(B)**. Correlation matrix for female PG glucose uptake. **(C)**. Correlation matrix for male ING glucose uptake. **(D)**. Correlation matrix for male PG glucose uptake.

